# The structure of mitochondrial genomes is associated with geography in *Arabidopsis thaliana*

**DOI:** 10.1101/2025.01.11.632530

**Authors:** Wenfei Xian, Zhigui Bao, Sebastian Vorbrugg, Yueqi Tao, Andrea Movilli, Ilja Bezrukov, Detlef Weigel

## Abstract

Chloroplasts and mitochondria are the primary sites for photosynthesis and respiration, each harboring its own unique genome. Although the organellar genomes are considerably smaller compared to the nuclear genome, they are nonetheless essential for survival of the organism. A common feature of many chloroplast and mitochondrial genomes is the presence of large repeated sequences longer than 1 kb. These can be either in inverted or direct orientation, and recombination between them leads to structural heteroplasmy. To understand the intraspecific evolution of organellar genomes, we assembled chloroplast and mitochondrial genomes of 143 *A. thaliana* accessions from PacBio HiFi sequencing data. We find large repeats to be associated with heteroplasmy and structural variation. Our extensive genome annotation identifies novel open reading frames (ORFs) in those accessions that lost large repeats, potentially introduced via horizontal gene transfer, illuminating additional paths for diversification of plant organelles. The loss of large repeats correlates with geography and phenotypes, pointing to their adaptive importance. The assembled and annotated organellar genomes constitute a rich source for future functional studies of the interaction between the three genomes of a plant.

## Introduction

Climate change-induced increases in temperature and drought affect plant photosynthesis and respiration (Ribas-Carbo et al. 2005; Hu et al. 2020; Trösch et al. 2022), threatening food security (Hasegawa et al. 2018). Photosynthesis occurs in chloroplasts, while respiration takes place in mitochondria. Both chloroplasts and mitochondria have their own independent genomes that have co-evolved closely with the nuclear genome (Sloan et al. 2018a; Lian et al. 2024b). While population-level genomic studies have deepened our understanding of species evolution and helped pinpoint alleles relevant to adaptation of wild and crop plants (1001 Genomes Consortium 2016; Wang et al. 2018; Liu et al. 2020; Zhou et al. 2022; Wilson et al. 2019; Evans et al. 2014; Jayakodi et al. 2020; Cheng et al. 2024), these studies have typically focused on the nuclear genome, with variation in the genomes of chloroplasts and mitochondria remaining less well understood. Of the efforts that have analyzed variation in organellar genomes at the population scale, these have mostly concentrated on chloroplast genomes (Go et al. 2024; Magdy et al. 2019), with analyses of mitochondrial genomes typically being based on a small number of assemblies (Sun et al. 2022; Wang et al. 2022; Fan et al. 2022).

In vascular plants, most chloroplast and mitochondrial genomes feature large repeated sequences longer than 1 kb (Wynn and Christensen 2019; Sloan 2013). Pairs of large repeats in either inverted or direct orientation have been found to trigger recombination, which leads in turn to structural heteroplasmy, defined as the presence of multiple chloroplast or mitochondrial genomes in a single individual (Palmer 1983; Kozik et al. 2019; Klein et al. 1994; Wang and Lanfear 2019). Heteroplasmy complicates genome representation, as a single linear sequence cannot encapsulate all configurations (Palmer 1983).

Frequent recombination-driven rearrangements hinder comparisons both between and within species, as whole-genome alignment software is primarily designed for conserved synteny (Li 2018). Rearrangements are biologically relevant, as they can impact fitness (Juszczuk et al. 2007; Shedge et al. 2010). Chimeric open reading frames (ORFs) resulting from rearrangements may cause cytoplasmic male sterility (Sandhu et al. 2007).

The chloroplast genome of the *Arabidopsis thaliana* reference accession Col-0 has a pair of inverted repeats that are 26 kb long (Sato et al. 1999). The mitochondrial genome of this accession contains two large repeats, the direct 4.2 kb long repeat I and the inverted 6.5 kb long repeat II (Unseld et al. 1997). Recombination between the two copies of the direct repeat I leads to two subgenomic DNAs of 134 and 234 kb (Klein et al. 1994; Unseld et al. 1997). The assembly of a small number of *A. thaliana* mitochondrial genomes from short or long reads has also shown that not all of them contain two copies of repeat II (Davila et al. 2011; Zou et al. 2022).

For this study, we assembled organellar genomes of 143 *A. thaliana* accessions from Pacbio HiFi data. We describe the structural diversity of both chloroplast and mitochondrial genomes, report a likely instance of horizontal gene transfer from another member of the Brassicaceae to *A. thaliana* mitochondrial genomes, and uncover an association between the loss of large repeats in mitochondrial genomes with geography and phenotypes.

## Results

### Structural Diversity of Organellar Genomes

We obtained PacBio HiFi reads for 143 *A. thaliana* accessions, for which nuclear genome assemblies have been recently reported (Lian et al. 2024a; Wlodzimierz et al. 2023; Kang et al. 2023; Rabanal et al. 2022). We used the TIPPo tool for assembly of chloroplast genomes (Xian et al. 2024). We used both TIPPo (Xian et al. 2024) and PMAT (Bi et al. 2024) to assemble mitochondrial genomes, manually selecting the completeness assembly for each accession. For three accessions (Geg-14, HR-10, Nz-1), assembly of the mitochondrial genome failed, and we used Verkko (Rautiainen et al. 2023) to perform whole-genome assemblies, visualized the assembly graph in Bandage (Wick et al. 2015). We then manually selected the appropriate nodes based on the structure of the subgraph, provided these as input to Ribotin (Rautiainen 2024) to extract the constituent HiFi reads, and finally reassembled these with flye (Kolmogorov et al. 2019).

Chloroplast genomes are found in the canonical form in all accessions, with one large single copy (LSC), one small single copy (SSC) and a pair of inverted repeats (IRs) (**Figure 1A** and **Figure S1**). The average size of the chloroplast genomes across all accessions is 154,297 bp, with a range of 152,495 bp to 154,909 bp (**Figure 1B** and **Table S1**).

**Figure 1.**
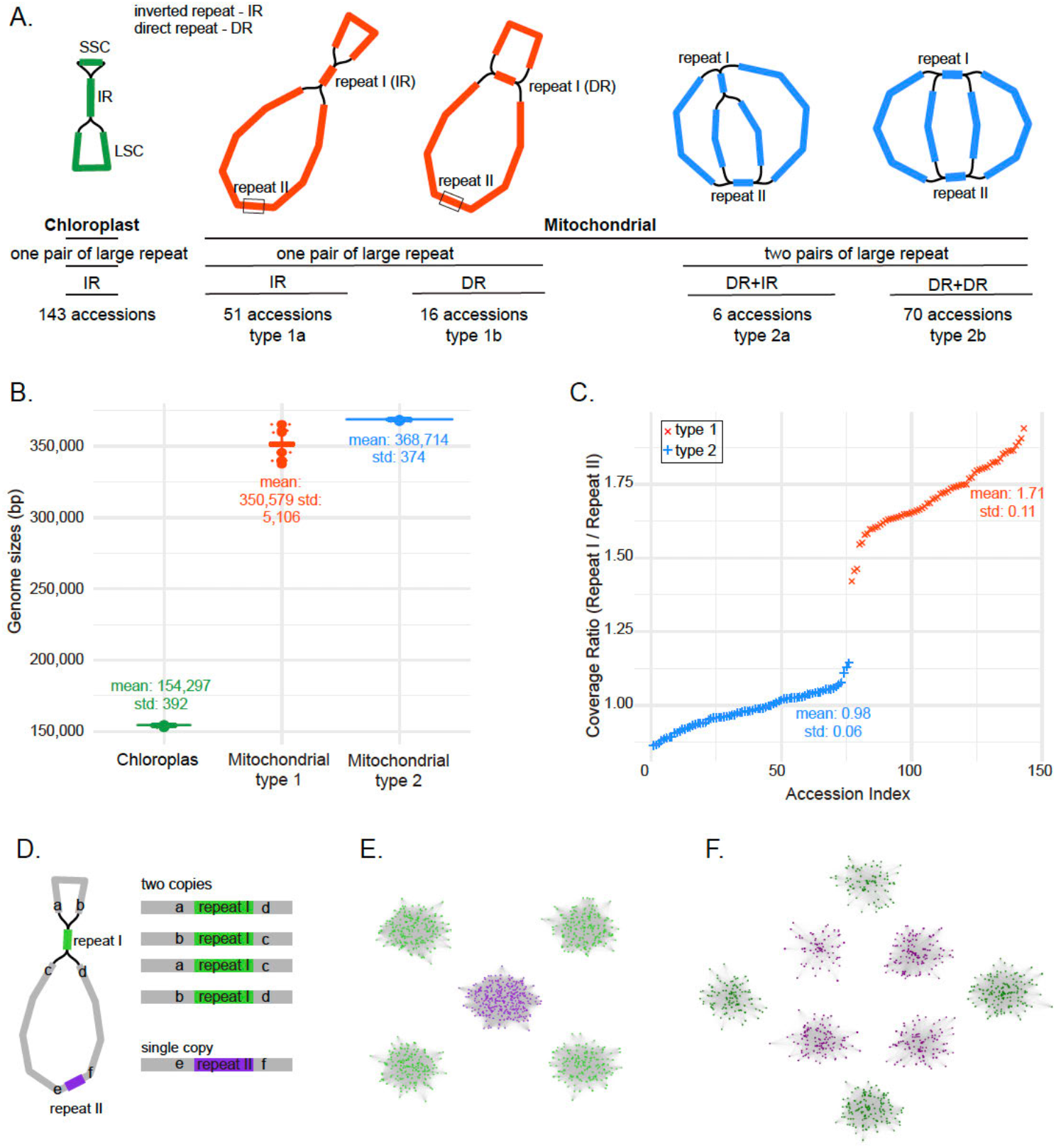
Structure diversity of organellar genomes. **A)** Major types of assembly graphs of chloroplast and mitochondrial genomes in *A. thaliana*. **B)** Distribution of genome sizes of chloroplast and mitochondrial genomes. **C)** Coverage ratio between repeat I and repeat II for inferring their relative copy number in mitochondrial genomes. **D)** Diagram illustrating the outcome of recombination with a pair of large repeats in a mitochondrial genome. **E)** Clusters of reads containing repeat I (green) and II (purple) sequences in accession Ler-0 (type 1). **F)** Clusters of reads containing repeat I (green) and II (purple) sequences in the reference accession Col-0 (type 2).

The mitochondrial genomes can be categorized into four primary types, distinguished by the copy number and orientation of large repeats within the assembly graph: 51 accessions of type 1a contain a pair of inverted repeat I and a single copy of repeat II; 16 accessions of type 1b have a pair of direct repeat I and a single copy of repeat II; 6 accessions of type 2a feature a pair of repeat I in direct orientation and a pair of repeat II in inverted orientation; and 70 accessions of type 2b have a pair of the two repeats, with both repeat I and II in inverted orientations (**Figure 1A, Figure S2** and **Table S2**). For accessions with two pairs of large repeats (types 2), the average mitochondrial genome size is 368,714 bp, ranging from 367,334 bp to 368,974 bp (**Figure 1B** and **Table S1**). The average mitochondrial genome size of accessions with only one pair of large repeats (type 1) is 350,579 bp and has a larger range, from 337,339 bp to 365,358 bp (**Figure 1B** and **Table S1**). The 67 mitochondrial genomes of type 1 are much more variable in size, with a standard deviation of 5,106 bp compared to 374 bp for the 76 type 2 genomes (**Figure 1B** and **Table S1**).

To validate the single copy of repeat II sequence in 67 accessions (type 1), we used read coverage to infer the copy number by mapping HiFi reads to repeat I and II sequences (**Table S3**). If the mitochondrial genome contains two copies of both repeat I and II (type 2), the coverage ratio of repeat I and repeat II should be close to 1. With two copies of repeat I and one copy of repeat II (types 1), the ratio should be closer to 2.

As expected, the mean coverage ratio for two copies of repeat I and II accessions (type 2) is 0.98 with a standard deviation of 0.06. However, the average coverage ratio for two copies of repeat I and one copy of repeat II accessions (type 2) is 1.71, with a standard deviation of 0.11 (**Figure 1C**). This did not change much when we considered only the first 4.2 kb of repeat II, which is over 50% longer than repeat I (**Table S4**). The variance to the expected ratio of 2 between repeat I and II in type 1 mitochondrial genomes cases suggests that these genomes have fewer copies of repeat I or more copies of repeat II than expected.

One possible explanation is the coexistence of type 2 mitochondrial genomes in these accessions, increasing the abundance of repeat II. A pair of large repeats has high recombination activity, resulting in heteroplasmy (Ramsey and Mandel 2019; Palmer 1983; Wang and Lanfear 2019; Sloan 2013). Assuming that repeat II has two copies, recombination will result in four combinations of repeat II and adjacent sequences, but only one combination when repeat II is present in only one copy (**Figure 1D**).

We searched for HiFi reads completely encompassing repeat II, and then extracted 1 kb sequences upstream and downstream of repeat II, and subjected these to an all-versus-all alignment using minimap2. Pairs of reads with > 95% mutual coverage were then provided as input for MCL clustering. As a control, the same analysis was conducted for repeat I. We found that in type 1 mitochondrial genome, repeat II reads formed only one cluster in each accession, indicating a lack of recombination (**Figure 1E, Table S5 and S6**). However, both repeat I of type 1 mitochondrial genome and repeats I and II of type 2 mitochondrial genomes formed four clusters in an accession, indicative of recombination activity (**Figure 1F, Table S5 and S6**). Thus, there is no evidence for the co-existence of the two different structural types of mitochondrial genomes in the type 1 accessions.

### Rearrangements in organellar genomes

Considering that recombination mediated by repeats can lead to rearrangements of organellar genomes (Davila et al. 2011; Cole et al. 2018). If we assume that each organellar genome contains only one pair of large repeats, two linear DNA genomes could result akin to haplotypes for individuals with a diploid nuclear genome (Wang and Lanfear 2019). Previous studies have often erroneously interpreted the chloroplast SSC as a hotspot for inversions that distinguish species because they generally ignored heteroplasmy caused by large repeats within individuals (Walker et al. 2015). This has happened as a consequence of the fact that prevailing alignment methods are built for comparison of linear DNA sequences. To avoid the impact of intragenomic recombination-induced rearrangements on alignments across individuals, we focus on rearrangements between single-copy regions and we do not use representations that connect the single-copy fragments into larger linear molecules via large repeats.

In this way, we identified rearrangement events through pairwise genome comparisons. If no rearrangement was detected, the genomes were classified as belonging to the same cluster. For each cluster, we randomly selected one accession as a representative.

We did not detect obvious rearrangement in chloroplast genomes (**Figure S3**). In mitochondrial genomes, however, rearrangements are frequent. In total, we identified 25 clusters (**Figure 2A, Figure S4**). The largest cluster, C1, contains 37 accessions (**Figure 2B**). When we projected these 25 clusters onto a nuclear phylogenetic tree (**Figure 2C**), we found that accessions belonging to the same cluster were not necessarily closely related, especially for larger clusters.

**Figure 2.**
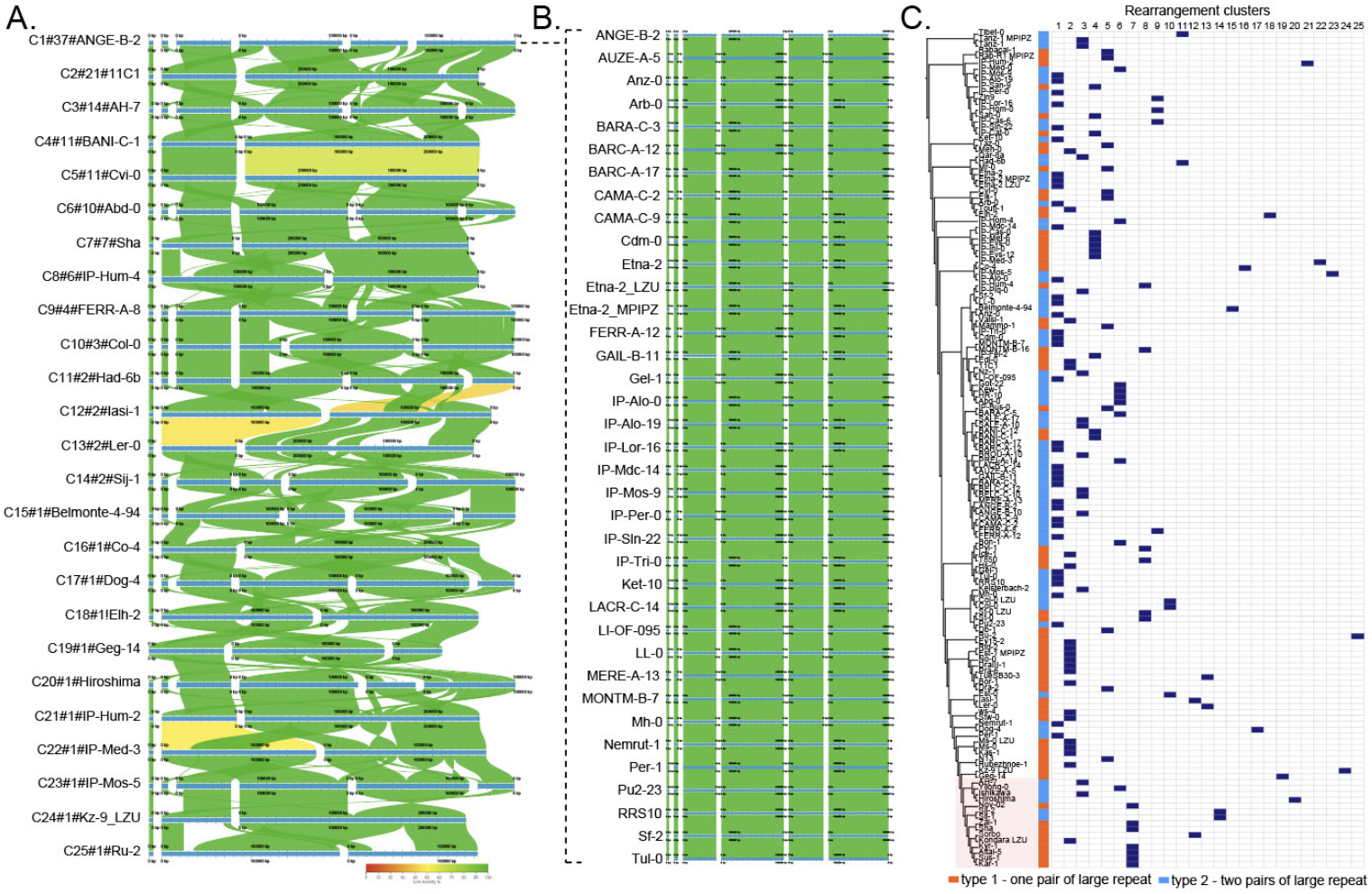
Rearrangements in mitochondrial Genomes. **A)** Whole genome alignments among the 25 clusters of mitochondrial genomes. Using “#” as a separator, the first string represents the cluster name, the second string indicates the number of accessions in the cluster, and the third string denotes the representative accession. **B)** Whole genome alignments of cluster 1. **C)** The distribution of cluster members across the nuclear phylogenetic tree.

### Evidence for horizontal gene transfer in mitochondrial genomes

Initially, we identified all ORFs of at least 50 amino acids across organellar genomes for 143 accessions and derived orthologous groups (OGs) with OrthoFinder (Emms and Kelly 2019). In the mitochondrial genome, rearrangements are considered as one of the sources for new ORFs (Wang et al. 2024). Since we found no rearrangements between chloroplast genomes, we did not expect to find many new ORFs in chloroplast genomes. Consistent with expectations, we identified 92 chloroplast OGs, of which 89 were conserved across all genomes. The remaining three, which were not similar to known proteins, were found in only one or two accessions, and we cannot exclude that these are due to assembly or annotation errors.

In the much larger mitochondrial genomes, we identified 625 OGs, with 516 missing in no more than one accession and 109 missing in at least two accessions (**Table S7**). Notably these 109 non-conserved ORFs were typically quite common. If the variation in ORFs is related to genome rearrangements, the distribution of OGs should correspond to the rearrangement-based clusters identified above (**Figure 2A**). We found that Presence/Absence Variation (PAV) of non-conserved OGs largely follow cluster assignments (**Figure 3A**). As absences are rarer than presences, we conclude that rearrangements are more likely to break existing ORFs than leading to the formation of new ORFs.

**Figure 3.**
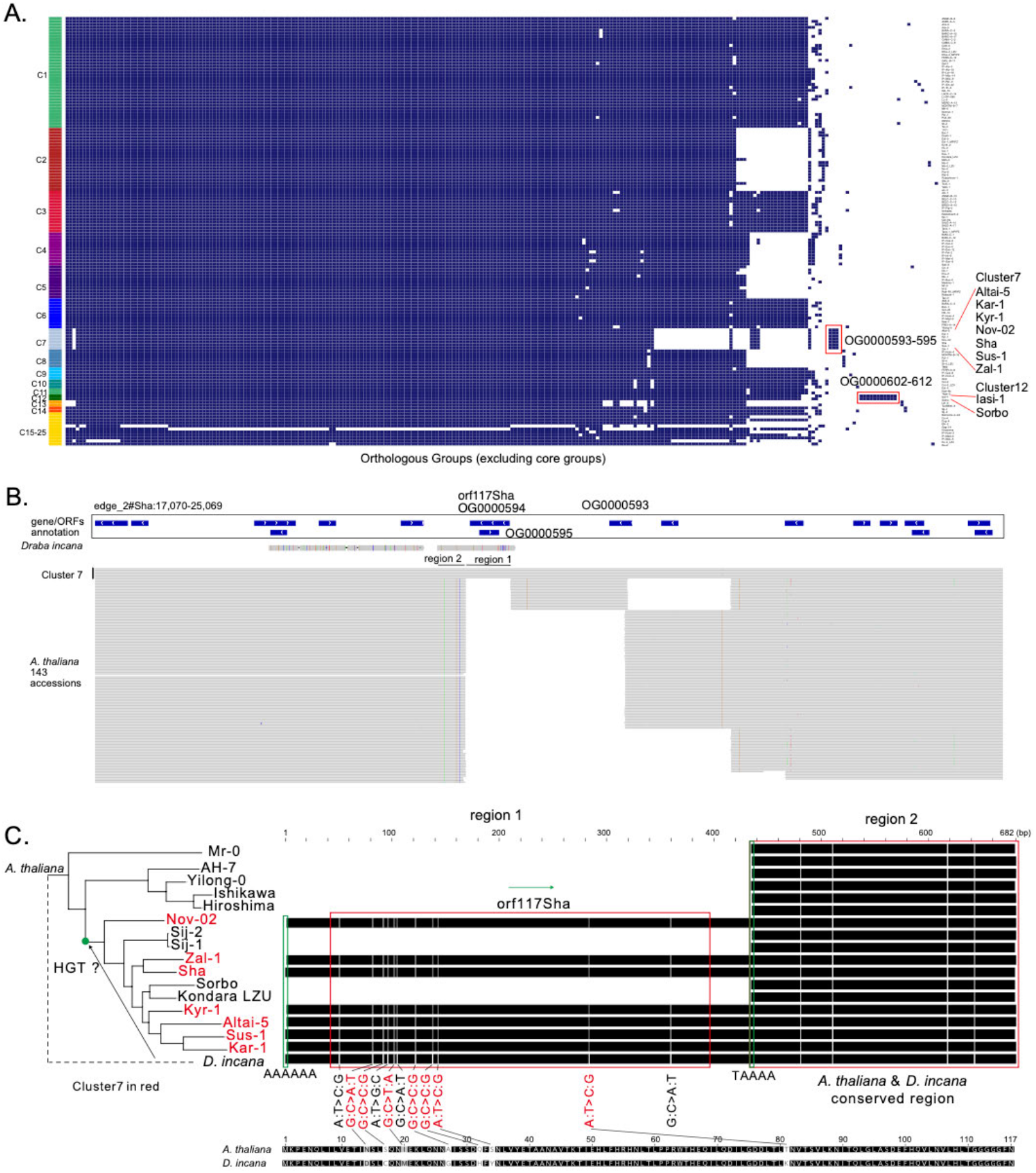
New ORFs in Mitochondrial genomes. **A)** Orthologous groups (OGs) of mitochondrial genes and ORFs. Accessions were ordered by their structures. **B)** IGV screenshot of the coverage of orf117Sha-related sequences in other accessions and annotation in the *D. incana* mitochondrial genome. Only one representative was chosen for each *A. thaliana* cluster. **C)** Close-up of the alignment of orf117Sha and adjacent sequences in several *A. thaliana* accessions.

We then focused on those OGs that are cluster-specific (cluster size ≥2). We found that a block with the three OGs OG0000593-595 is specific to cluster 7, and that a block with the 11 OGs OG0000602-612 is specific to cluster 12. For each OG, we selected a representative sequence and aligned it using DIAMOND BLASTP (Buchfink et al. 2021) to the UniRef90 database, where 320 OGs matched homologous sequences under thresholds of ≥75% coverage and ≥75% identity (**Figure S5**). OG000608, unique to cluster 12, aligned with a *Coffea canephora* protein from unassembled WGS sequence at close to 100% coverage but only about 30% amino acid identity (**Figure S5**). Specifically, the homologous sequences for OG000594, which is unique to cluster 7, showed 100% coverage and identity in all seven accessions of this cluster. This ORF was previously reported as orf117Sha in *A. thaliana* accession Sha, where it is associated with cytoplasmic male sterility (CMS) (Gobron et al. 2013). The original report also indicated that this ORF was detectable in accessions from Central Asia, which agrees with our finding that cluster 7 is restricted to this geographic region (Lian et al. 2024a).

It had been suggested that orf117Sha originates from an unknown source (Gobron et al. 2013). Given that OGs OG000593-595 are specific to cluster 7, it is likely that the sequences containing the ORFs in these OGs are unique to cluster 7 at the genomic level – indeed, our dataset shows the orf117Sha region to be unique to cluster 7 (**Figure 3A and 3B**). Searching the NT database, we found a remarkably close hit, 96.8% at the DNA level, in the mitochondrial genome of another Brassicaceae species, *Draba incana* (GenBank: OY755218.1, data from Darwin Tree of Life project), as well as more distant hits in the mitochondria of other plants, including a member of the Fabaceae (**Figure S6**).

With a larger region from *D. incana* around the orf117Sha related sequence as a query, we identified homologous sequences in five closely related *A. thaliana* accessions in addition to the OG000594 region in cluster 7 accessions (**Figure 3C**). Based on a multiple sequence alignment, the *D. incana* orf117Sha related sequence block can be divided into two regions: Region 1, which includes orf117Sha plus about 30 bp on either side and which is present only in all cluster 7 accessions; region 2 is conserved between *D. incana* and *A. thaliana* (**Figure 3B & 3C**).

Comparing Region 1/orf117Sha between *A. thaliana* and *D. incana*, we find 11 single-nucleotide differences, seven of which are non-synonymous mutations. The adjacent Region 2, which is only about half as long as Region 1, contains only four single-nucleotide differences. Additionally, we detected adenine (A)-rich micro-homologies enriched at both ends of Region 1/orf117Sha (**Figure 3C**). The seven accessions of cluster 7 and four additional accessions Sij-2, Sij-1, Sorbo and Kondara_LZU form a monophyletic clade, as can be inferred from Figure 3C, consistent with a potential HGT event when this clade arose.

For the 11 cluster 12-specific OGs (3,427 bp), we also observed a potential origin from HGT. The best match in the NT database belongs to the mitochondrial genome of *Arabidopsis lyrata* (NC_081483.1, coverage 99%, similarity 99.33%), for which no further provenance information is available (**Figure S7**). We assembled the complete mitochondrial genomes of *A. lyrata* accessions MN47 and NT1 (Wlodzimierz et al. 2023) using PacBio HiFi reads, but did not find any homologous sequences, suggesting that this cluster also segregates in *A. lyrata*.

### Loss of repeat II is associated with geography

We analyzed the distribution of repeat I and II in mitochondrial genomes on a larger scale using published Illumina short read sequencing data of 995 accessions (**Table S8**) (1001 Genomes Consortium 2016; Zou et al. 2017). We first confirmed that short read coverage can be used to infer the relative copy number with 38 accessions that had sequenced with both Pacbio HiFi long reads and Illumina short reads, ensuring that the read sets were indeed from the same accessions (**Figure S8**). Coverage ratios were highly consistent between long and short reads (**Figure S9**).

We observed that HPG1 accessions, members of a nearly clonal lineage that was introduced to North America about 400 years ago (Exposito-Alonso et al. 2018), contain two copies of repeat I and repeat II, indicating that the copy number of repeat II has not changed over approximately 400 years of evolution **(Figure S10**). In Central Asia, accessions with only one pair of large repeats (type 1) predominate, but in the Yangtze River Basin (Zou et al. 2017) are mostly accessions with two pairs of large repeats (types 2) (**Figure 4A**).

**Figure 4.**
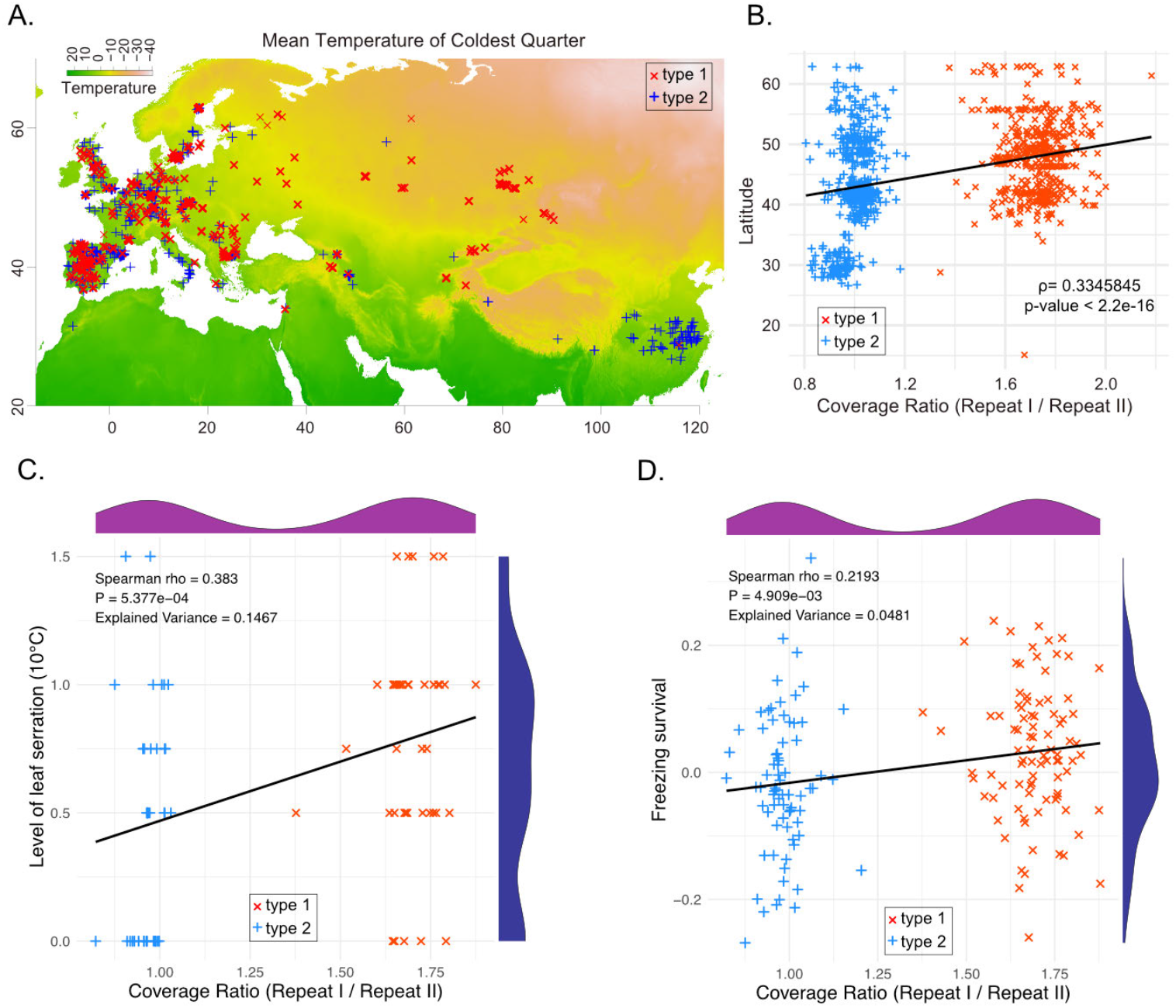
Loss of large repeats is associated with local geography and potentially local adaptation. **A)** Geographic distribution of 995 accessions in which repeat structure of mitochondrial genomes was estimated with short reads. **B)** Correlation between coverage ratio and latitude. Spearman’s ρ and P-values are shown. **C)** Correlation between coverage ratio and level of leaf serration in 10°C. **D)** Correlation between coverage ratio and freezing survival.

Turning to geography more broadly, there was a significant positive correlation between latitude and coverage ratio (ρ = 0.33, p-value < 2.2e-16), indicating that accessions with two pairs of repeats are more common at lower latitudes, while those with one pair of repeats are more common at higher latitudes (**Figure 4B**). There was no significant correlation with longitude (p-value = 0.099) (**Figure S11**).

We examined whether the loss of repeat II is associated with specific phenotypes. We performed a correlation analysis with 1,695 phenotypes (Voichek and Weigel 2020); only 13 phenotypes showed a correlation that was significant at a p-value threshold of 0.01 (**Table S9**). The strongest correlation was with leaf serration at 10°C (p-value = 0.0005, rho = 0.383) (**Figure 4C**), with higher phenotypic values corresponding to sharper and more jagged serrations (Atwell et al. 2010). The leaves of type 1 accessions are more highly serrated than those of type 2 accessions in 10°C. However, differences in leaf serration at 16°C and 22°C were not statistically significant, suggesting a connection to cold adaptation. One other significant phenotype was also related to cold adaptation: freezing survival (p-value = 0.005, rho = 0.219) (Horton et al. 2016), with type I accessions having higher survival rates upon freezing.

## Discussion

Pioneering research conducted 40 years ago in seven angiosperm species revealed extensive rearrangements in the chloroplast genomes of two legume species, pea and broad bean, both of which are characterized by the loss of a large inverted repeat sequence (Palmer and Thompson 1982). In contrast, virtual collinearity is observed among the chloroplast genomes of spinach, petunia, and cucumber, all of which retain both copies of the inverted repeat (Palmer and Thompson 1982). This finding suggested that genomes containing inverted repeats tend to be relatively more stable, whereas genomes that have lost the large repeats are more dynamic. Similar patterns of increased rearrangement rates were observed in the genus *Cephalotaxus*, which had also lost its inverted repeats (Ji et al. 2021).

Previous studies of inter- and intraspecific organellar genome variation in plants have primarily focused on plastid genomes. Here, we extend these studies through a large-scale investigation of large repeats in mitochondrial genomes of *A. thaliana* accessions. We confirm that genomes with two copies of repeat I and II are more stable, whereas those that have lost one copy of repeat II are more diverse. Specifically, mitochondrial genome size is stable in accessions with two copies, but it becomes more variable in those with only one copy of repeat II (**Figure 1B**). Furthermore, based on the PAV distribution of ORFs (OGs), the mitochondrial genomes of accessions in clusters 1, 3, and 6, which contain two copies of the repeats, are similar, whereas mitochondrial genomes in clusters 2, 4, 5, 7, and 8, which contain only one copy of repeat II, show significant differences (**Figure 3A**).

Heteroplasmy in plant organellar genomes is an intriguing phenomenon that poses several analytical challenges. Heteroplasmy complicates the representation of organellar genomes, which cannot be simply depicted as a single linear DNA sequence. Although the sequence content may remain unchanged, the sequence order can vary, making it difficult to identify structural variations across different species or even within different individuals of the same species, potentially leading to misinterpretation (Walker et al. 2015).

We wanted to minimize the impact of large repeat-mediated heteroplasmy—which can result in multiple genome sequences with identical content but different arrangements. We therefore did not connect non-repetitive fragments through large repeats into a single master ring. Instead, we directly compared the rearrangements between each fragment (**Figure 3A**). This approach is similar to a recent method used for identifying structural variations (SVs) in the *A. thaliana* nuclear genome, where the genome is divided into chunks before alignment to a reference genome, in order to avoid alignment errors due to rearrangements (Igolkina et al. 2024). The same principle can be applied to transposon alignments (Anderson et al. 2019), as orthologous transposons typically do not reside within large orthologous blocks in whole-genome alignments.

With the sequencing of more and more genomes, reports of horizontal gene transfer are increasing (Haimlich et al. 2024; Cheng et al. 2019). In this study, the *orf117Sha* gene, which was thought to have come from an “unknown source” (Gobron et al. 2013) due to the limited number of sequenced genomes at the time, can now be linked to a very similar sequence found in another Brassicaceae, *D. incana* (Bright 2022), with the majority of other Brassicaceae lacking orf117Sha related sequences. Given that the *A. thaliana* accessions with *orf117Sha* form a monophyletic clade in a nuclear phylogenetic tree and that the orf117Sha sequence is identical, if it was transferred by HGT, it likely took place in the ancestor of these accessions. A recent study revealed that the parasitic species *Lophophytum mirabile* acquires up to 74% of its mitochondrial DNA through recurrent horizontal gene transfer and the authors proposed that horizontally transferred mitochondrial DNA segments become circularized via microhomology-mediated repair pathways, subsequently integrating into the mitochondrial genome through recombination (Emilia Roulet et al. 2024). Similarly, in our findings, we observed A-rich repeats flanking the insertion fragments, consistent with the formation of circular DNA being an important ingredient of HGT of organellar DNA.

Although recent experimental work demonstrated that the removal of the inverted repeats from the tobacco plastid genome by genome editing had little impact on gross morphology or stress responses (Krämer et al., 2024), we found that in *A. thaliana*, the loss of large repeats in the mitochondrial genome is associated with geography and temperature-related phenotypes, in support of the evolutionary relevance of variation in organellar genomes.

## Materials and Methods

### Assembling organellar genomes

For each accession, we randomly selected PacBio HiFi reads corresponding to 1 and 2 Gb. For the chloroplast genomes, we used the assembly graph generated by TIPPo (Xian et al. 2024). For the mitochondrial genomes, we used both TIPPo v1.0 (Xian et al. 2024) and PMAT v1.5.3 (Bi et al. 2024) for primary assembly. We visualized each assembly graph using Bandage v0.9.0 (Wick et al. 2015) and manually selected the complete assembly graph. In three accessions (Geg-14, HR-10, Nz-1), we were unable to obtain a complete mitochondrial genome using TIPPo or PMAT. For these accessions, we first performed whole-genome assembly with Verkko v1.4.1 (Rautiainen et al. 2023) followed by visual inspection of the graph to identify mitochondrial nodes. We used Ribotin v1.2 (Rautiainen 2024) to extract the corresponding HiFi reads and finally assembled the mitochondrial genome using Flye 0.7.17-r1188 (Kolmogorov et al. 2019).

### Estimating the copy number of large repeats using sequencing depth

Considering that repeat I and repeat II are highly conserved in *A. thaliana* mitochondrial genomes, we aligned the PacBio HiFi data from each accession to the published *A. thaliana* Col-0 reference sequences of repeat I and repeat II (Sloan et al. 2018b) using minimap2 0.7.17-r1188 (Li 2018), retaining alignments with coverage of at least 90% of the length of the reference sequence. Next, we used mosdepth v0.3.3 (Pedersen and Quinlan 2018) to calculate the sequencing depth, and derived the ratio of sequencing depths of repeat I to repeat II. The same method was applied to Illumina 100 bp paired-end short-read sequencing data (1001 Genomes Consortium 2016; Zou et al. 2017), except that we used BWA 0.7.17-r1188 (Li and Durbin 2009) as aligner.

### Estimating the extent of heteroplasmy

To estimate the abundance of each heteroplasmic genome, we used GraphAligner v1.0.17 (Rautiainen and Marschall 2020) to align PacBio HiFi reads to the organellar assembly graph, ensuring that the reads fully spanned the large repeats. The relative orientation of the nodes adjacent to the large repeats was used to define different heteroplasmy states. Finally, we counted the number of reads corresponding to each state to determine their respective abundances.

### Detecting rearrangements

First, we performed pairwise whole genome alignments of mitochondrial genomes using minimap2 0.7.17-r1188 (Li 2018). In the absence of rearrangements, the alignment target coverage should be close to 100%, and the number of alignment blocks should equal the number of fragments in the target genome. Based on this principle, we first identified pairs of accessions without evidence of rearrangements. We then clustered mitochondrial genomes without apparent rearrangements with Markov Cluster Algorithm (MCL) (14-137) (Enright et al. 2002). We performed multiple sequence alignment of all genomes in each cluster using minitv ([CSL STYLE ERROR: reference with no printed form.]) and visualized the results with AliTV (Ankenbrand et al. 2017). Finally, we visually inspected the alignments to validate the absence of obvious errors in clustering.

### Construction of nuclear phylogenetic tree

The nuclear phylogenetic tree was constructed using Mashtree v1.4.6 (Katz et al. 2019), with the assembly of each accession as input.

### Annotation of Organellar Genomes

For each organellar genome, we identified all ORFs longer than 149 bp beginning with a start and ending with a stop codon with Getorf (Rice et al. 2000). We annotated the core genes using miniprot 0.12-r237 (Li 2023) based on the Col-0 mitochondrial genome annotation (Sloan et al. 2018b). We removed ORFs that overlapped with the core genes to generate the final annotation.

We used OrthoFinder v2.5.4 (Emms and Kelly 2019) to identify orthologous gene families. With the OrthoGroup (OG) results in hand, we selected a representative sequence for each OG based on median length of the OG and used DIAMOND v2.0.13 (Buchfink et al. 2015) BLASTp to align these sequences against the UniRef90 database. If a match was found under the default threshold, we selected the best alignment as the representative match.

## Data availability

HiFi reads used in this study were downloaded from ENA/NCBI/CNCB: PRJEB62038, PRJEB55353, PRJEB55632, PRJEB50694 and PRJCA012695 (Lian et al. 2024a; Wlodzimierz et al. 2023; Kang et al. 2023; Rabanal et al. 2022).

## Acknowledgments

We thank Wei Yuan, Li He and Haim Ashkenazy for the discussions. This study was supported by the Max Planck Society and the Novozymes Prize of the Novo Nordisk Foundation (D.W.).

