## Supplementary Figures for "The structure of mitochondrial genomes is associated with geography in *Arabidopsis thaliana*"

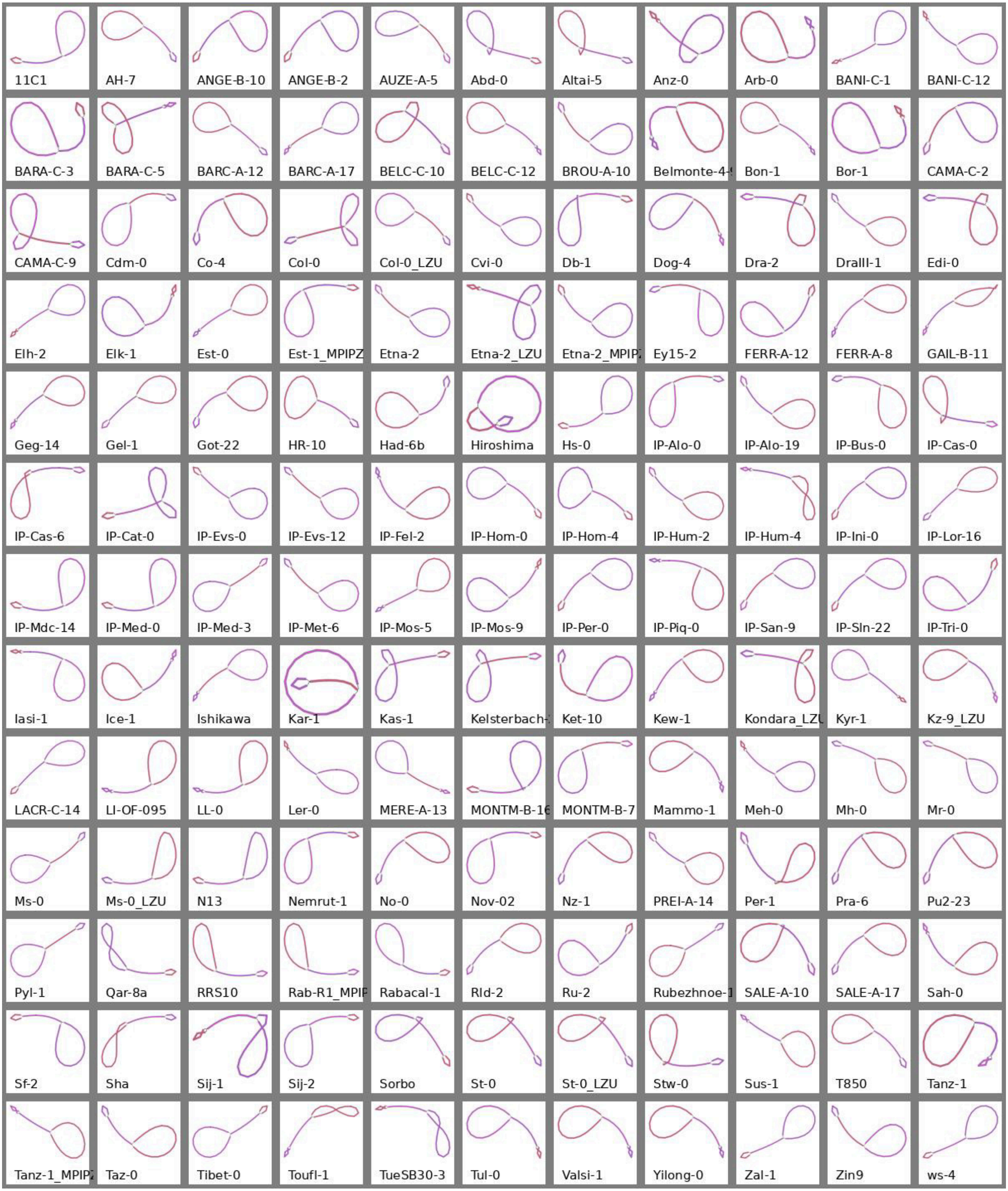


**Figure S1.** Visualization of chloroplast assembly graphs for 143 *Arabidopsis thaliana* accessions.

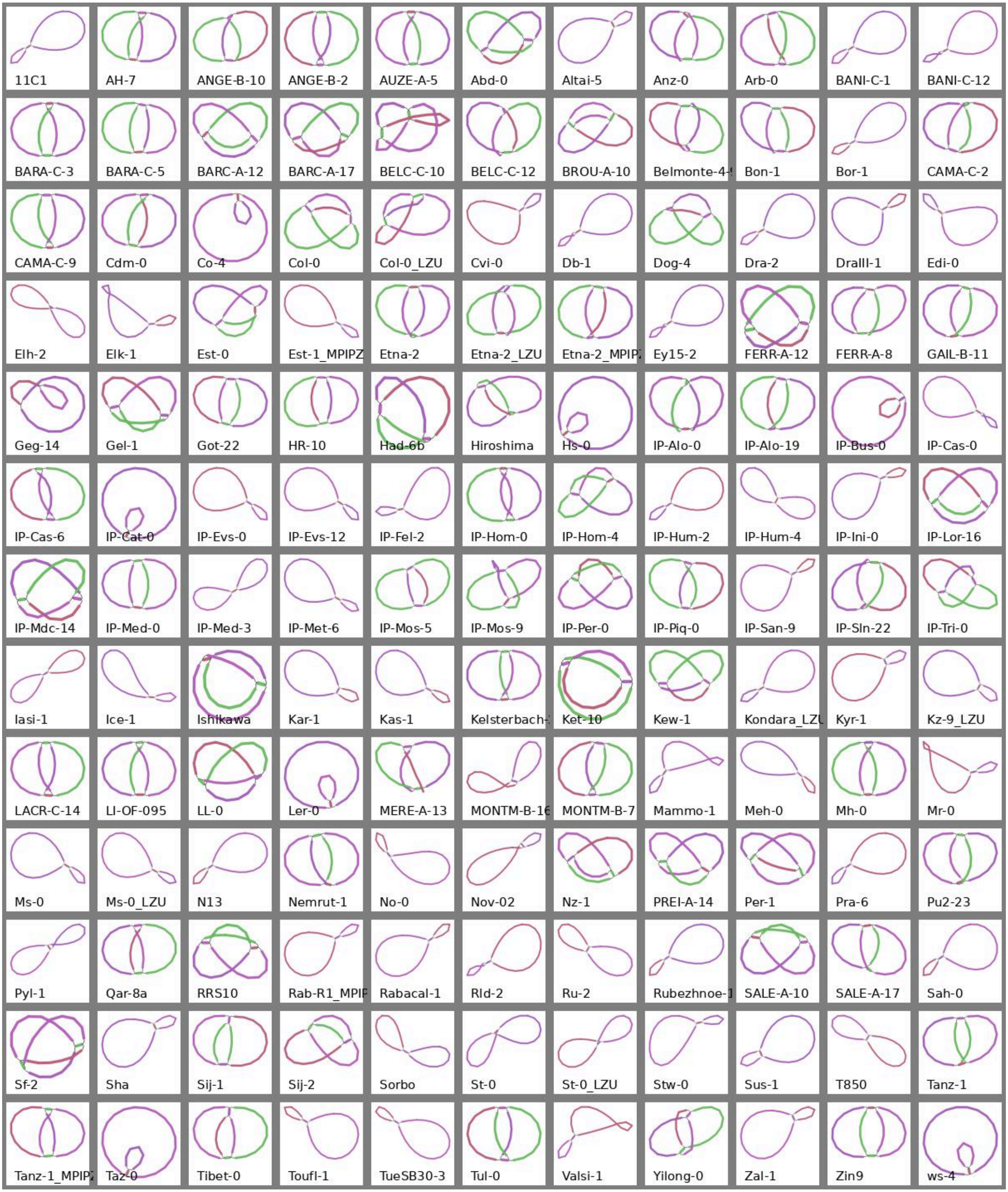


**Figure S2.** Visualization of mitochondrial assembly graphs for 143 *Arabidopsis thaliana* accessions.

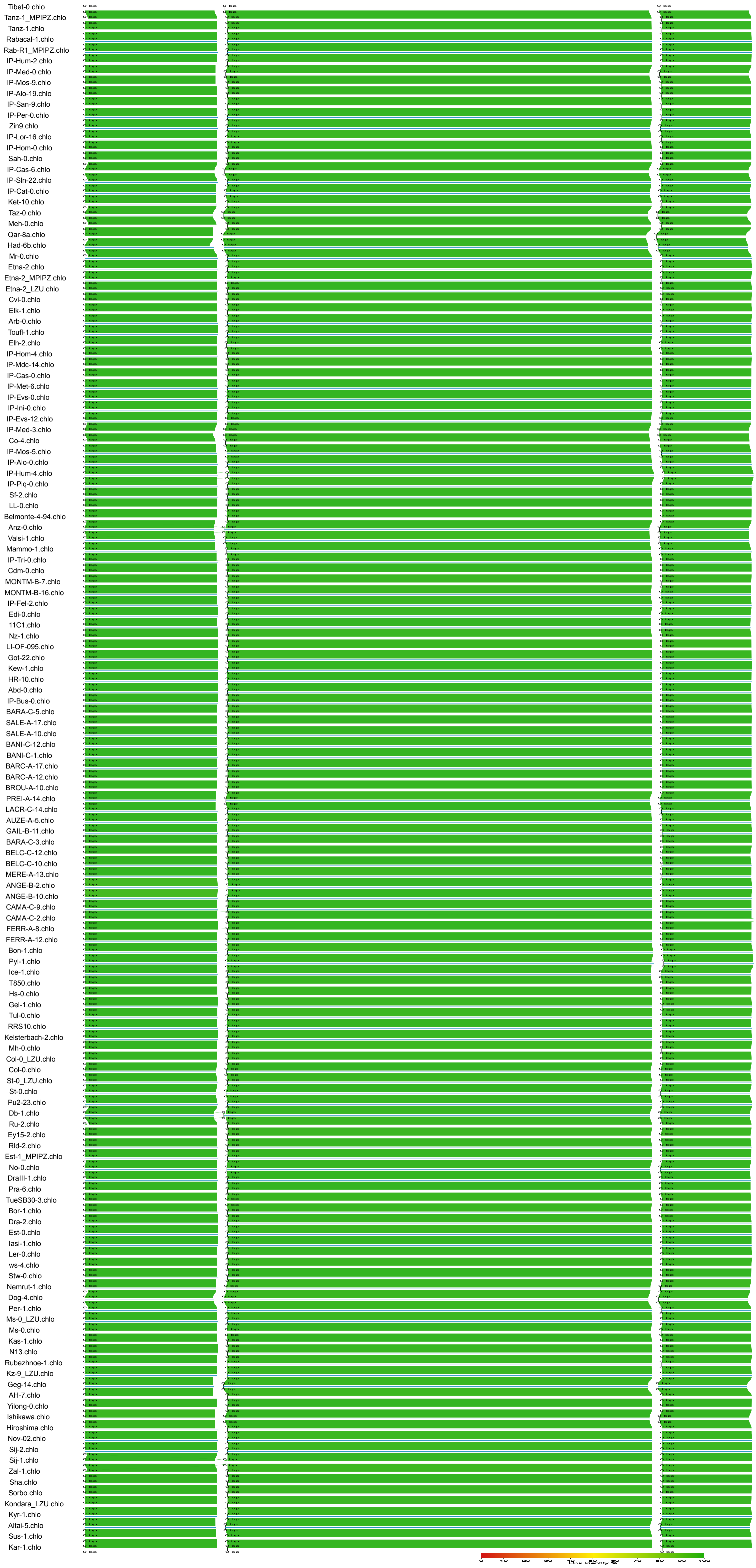


**Figure S3.** Multiple whole genome alignment of chloroplast genome among 143 accessions.

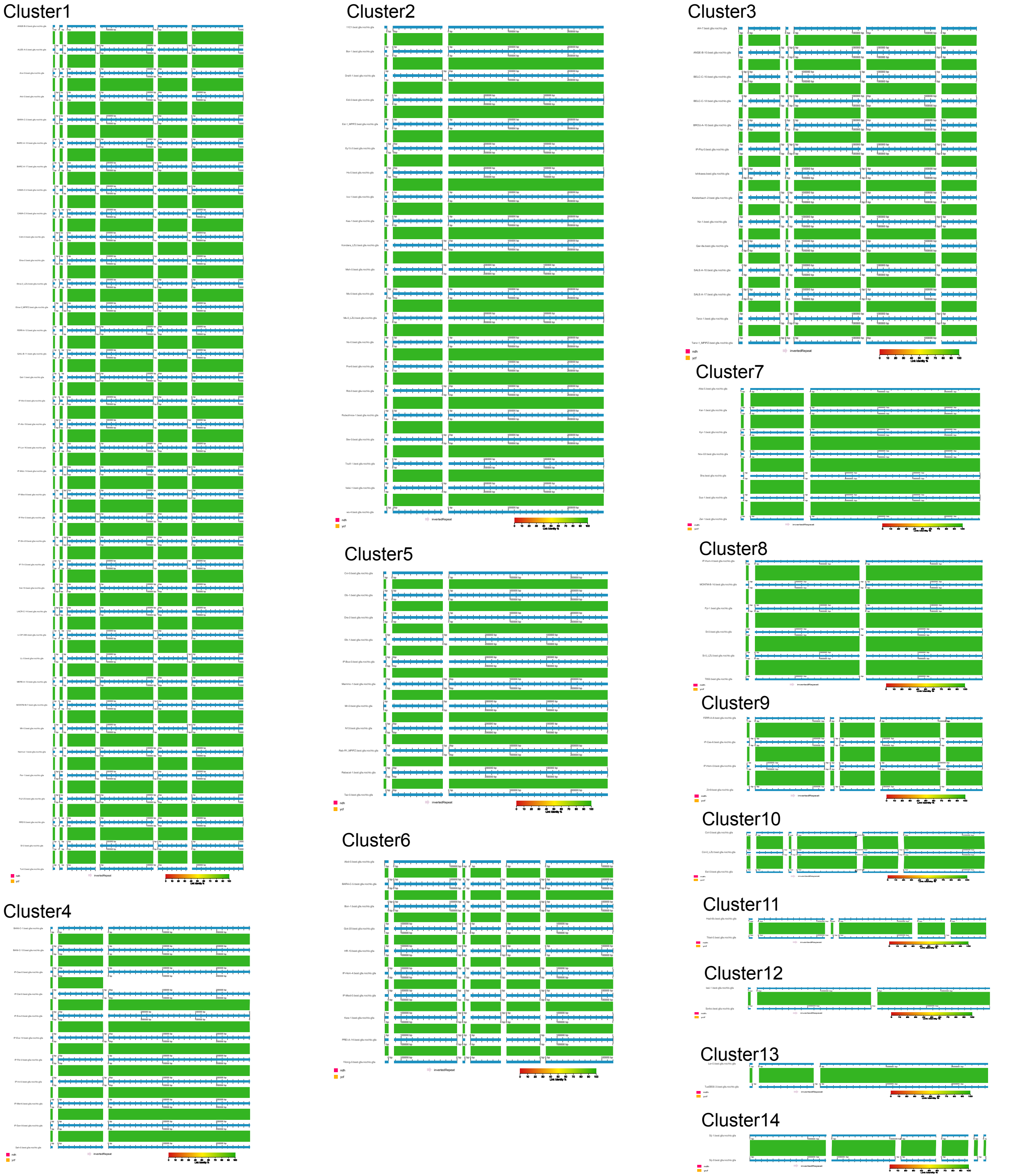


**Figure S4.** Mulitple / pair whole genome alignment for each mitochondrial genome cluster.

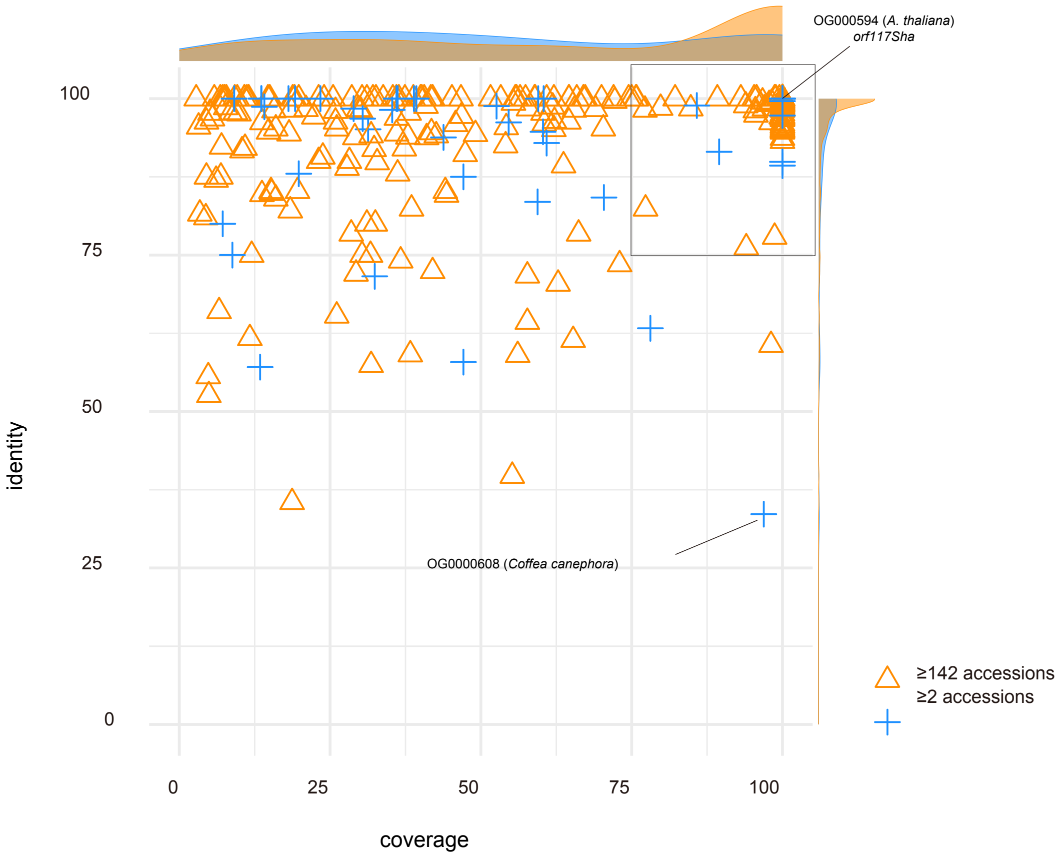


**Figure S5.** The alignment of representative sequences from mitochondrial genes orthologous groups to uniref90 database.

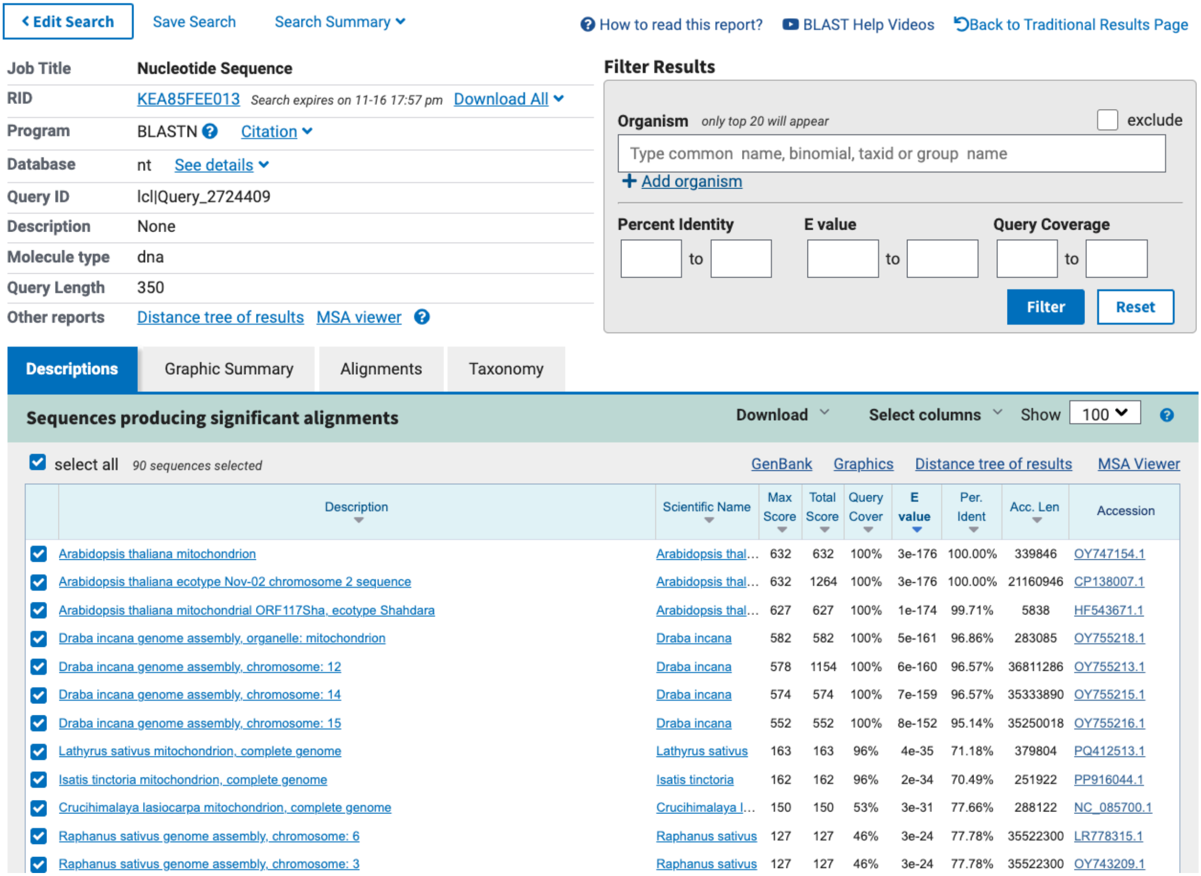

< Edit Search | Save Search | Search Summary | How to read this report? | BLAST Help Videos | Back to Traditional Results Page

| | |
|---|---|
| Job Title | Nucleotide Sequence |
| RID | KEA85FEE013 Search expires on 11-16 17:57 pm Download All |
| Program | BLASTN Citation |
| Database | nt See details |
| Query ID | lcl\|Query_2724409 |
| Description | None |
| Molecule type | dna |
| Query Length | 350 |
| Other reports | Distance tree of results MSA viewer |

Filter Results

Organism *only top 20 will appear* exclude

Type common name, binomial, taxid or group name

+ Add organism

Percent Identity to | E value to | Query Coverage to

Filter Reset

Descriptions | Graphic Summary | Alignments | Taxonomy

**Sequences producing significant alignments** Download Select columns Show 100

select all *90 sequences selected* GenBank Graphics Distance tree of results MSA Viewer

| Description | Scientific Name | Max Score | Total Score | Query Cover | E value | Per. Ident | Acc. Len | Accession |
|---|---|---|---|---|---|---|---|---|
| Arabidopsis thaliana mitochondrion | Arabidopsis thal... | 632 | 632 | 100% | 3e-176 | 100.00% | 339846 | OY747154.1 |
| Arabidopsis thaliana ecotype Nov-02 chromosome 2 sequence | Arabidopsis thal... | 632 | 1264 | 100% | 3e-176 | 100.00% | 21160946 | CP138007.1 |
| Arabidopsis thaliana mitochondrial ORF117Sha, ecotype Shahdara | Arabidopsis thal... | 627 | 627 | 100% | 1e-174 | 99.71% | 5838 | HF543671.1 |
| Draba incana genome assembly, organelle: mitochondrion | Draba incana | 582 | 582 | 100% | 5e-161 | 96.86% | 283085 | OY755218.1 |
| Draba incana genome assembly, chromosome: 12 | Draba incana | 578 | 1154 | 100% | 6e-160 | 96.57% | 36811286 | OY755213.1 |
| Draba incana genome assembly, chromosome: 14 | Draba incana | 574 | 574 | 100% | 7e-159 | 96.57% | 35333890 | OY755215.1 |
| Draba incana genome assembly, chromosome: 15 | Draba incana | 552 | 552 | 100% | 8e-152 | 95.14% | 35250018 | OY755216.1 |
| Lathyrus sativus mitochondrion, complete genome | Lathyrus sativus | 163 | 163 | 96% | 4e-35 | 71.18% | 379804 | PQ412513.1 |
| Isatis tinctoria mitochondrion, complete genome | Isatis tinctoria | 162 | 162 | 96% | 2e-34 | 70.49% | 251922 | PP916044.1 |
| Crucihimalaya lasiocarpa mitochondrion, complete genome | Crucihimalaya l... | 150 | 150 | 53% | 3e-31 | 77.66% | 288122 | NC_085700.1 |
| Raphanus sativus genome assembly, chromosome: 6 | Raphanus sativus | 127 | 127 | 46% | 3e-24 | 77.78% | 35522300 | LR778315.1 |
| Raphanus sativus genome assembly, chromosome: 3 | Raphanus sativus | 127 | 127 | 46% | 3e-24 | 77.78% | 35522300 | OY743209.1 |

**Figure S6.** The Sha orf117 sequence was aligned against the NCBI NT database using BLAST.

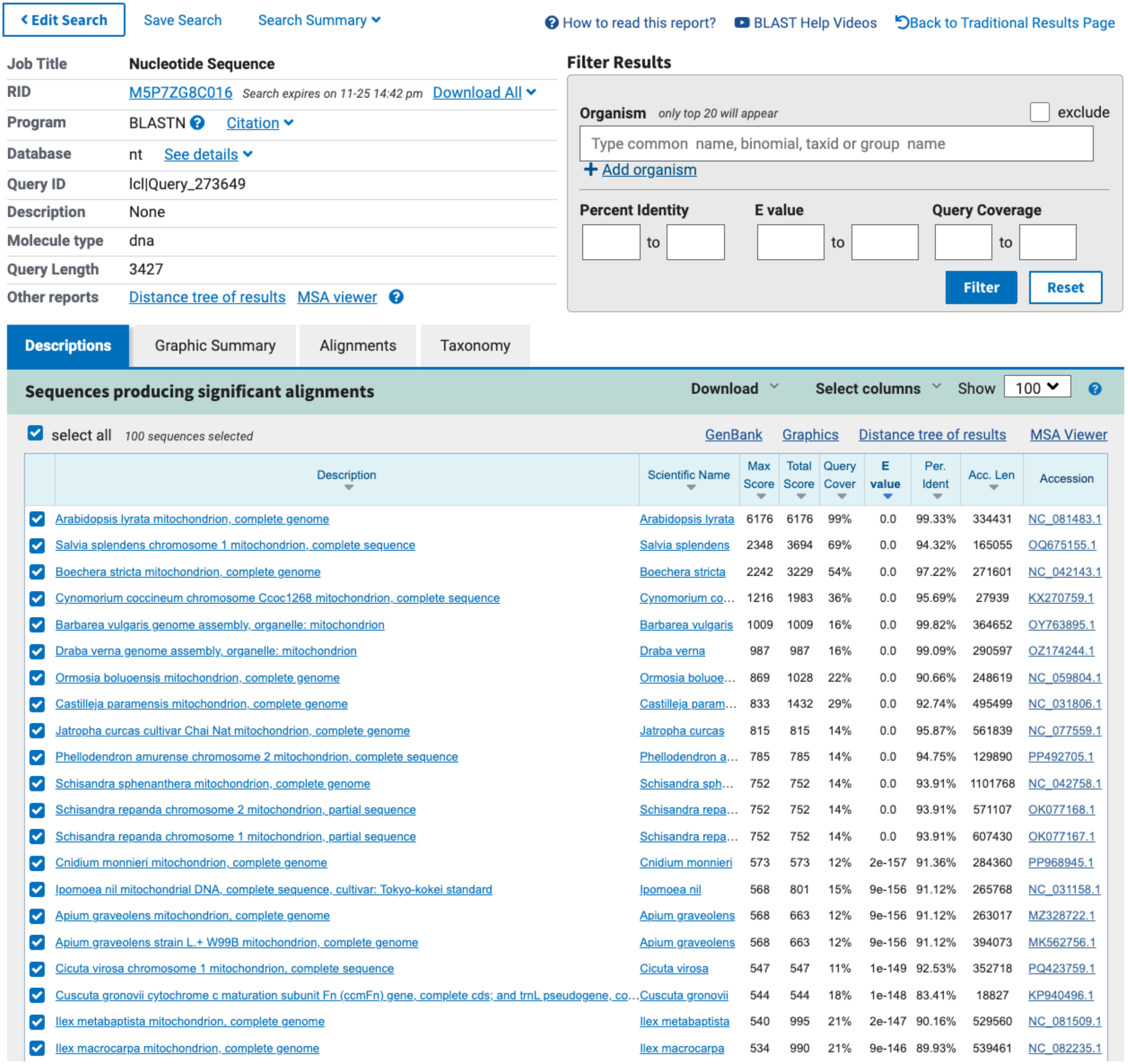

< Edit Search | Save Search | Search Summary | How to read this report? | BLAST Help Videos | Back to Traditional Results Page

| | |
|---|---|
| Job Title | Nucleotide Sequence |
| RID | M5P7ZG8C016 *Search expires on 11-25 14:42 pm* Download All |
| Program | BLASTN Citation |
| Database | nt See details |
| Query ID | lcl\|Query_273649 |
| Description | None |
| Molecule type | dna |
| Query Length | 3427 |
| Other reports | Distance tree of results MSA viewer |

**Filter Results**

Organism *only top 20 will appear* exclude

Type common name, binomial, taxid or group name

+ Add organism

Percent Identity to | E value to | Query Coverage to

Filter | Reset

Descriptions | Graphic Summary | Alignments | Taxonomy

**Sequences producing significant alignments** Download Select columns Show 100

select all *100 sequences selected* GenBank Graphics Distance tree of results MSA Viewer

| Description | Scientific Name | Max Score | Total Score | Query Cover | E value | Per. Ident | Acc. Len | Accession |
|---|---|---|---|---|---|---|---|---|
| Arabidopsis lyrata mitochondrion, complete genome | Arabidopsis lyrata | 6176 | 6176 | 99% | 0.0 | 99.33% | 334431 | NC_081483.1 |
| Salvia splendens chromosome 1 mitochondrion, complete sequence | Salvia splendens | 2348 | 3694 | 69% | 0.0 | 94.32% | 165055 | OQ675155.1 |
| Boechera stricta mitochondrion, complete genome | Boechera stricta | 2242 | 3229 | 54% | 0.0 | 97.22% | 271601 | NC_042143.1 |
| Cynomorium coccineum chromosome Ccoc1268 mitochondrion, complete sequence | Cynomorium co... | 1216 | 1983 | 36% | 0.0 | 95.69% | 27939 | KX270759.1 |
| Barbarea vulgaris genome assembly, organelle: mitochondrion | Barbarea vulgaris | 1009 | 1009 | 16% | 0.0 | 99.82% | 364652 | OY763895.1 |
| Draba verna genome assembly, organelle: mitochondrion | Draba verna | 987 | 987 | 16% | 0.0 | 99.09% | 290597 | OZ174244.1 |
| Ormosia boluoensis mitochondrion, complete genome | Ormosia boluoe... | 869 | 1028 | 22% | 0.0 | 90.66% | 248619 | NC_059804.1 |
| Castilleja paramensis mitochondrion, complete genome | Castilleja param... | 833 | 1432 | 29% | 0.0 | 92.74% | 495499 | NC_031806.1 |
| Jatropha curcas cultivar Chai Nat mitochondrion, complete genome | Jatropha curcas | 815 | 815 | 14% | 0.0 | 95.87% | 561839 | NC_077559.1 |
| Phellodendron amurense chromosome 2 mitochondrion, complete sequence | Phellodendron a... | 785 | 785 | 14% | 0.0 | 94.75% | 129890 | PP492705.1 |
| Schisandra sphenanthera mitochondrion, complete genome | Schisandra sph... | 752 | 752 | 14% | 0.0 | 93.91% | 1101768 | NC_042758.1 |
| Schisandra repanda chromosome 2 mitochondrion, partial sequence | Schisandra repa... | 752 | 752 | 14% | 0.0 | 93.91% | 571107 | OK077168.1 |
| Schisandra repanda chromosome 1 mitochondrion, partial sequence | Schisandra repa... | 752 | 752 | 14% | 0.0 | 93.91% | 607430 | OK077167.1 |
| Cnidium monnieri mitochondrion, complete genome | Cnidium monnieri | 573 | 573 | 12% | 2e-157 | 91.36% | 284360 | PP968945.1 |
| Ipomoea nil mitochondrial DNA, complete sequence, cultivar: Tokyo-kokei standard | Ipomoea nil | 568 | 801 | 15% | 9e-156 | 91.12% | 265768 | NC_031158.1 |
| Apium graveolens mitochondrion, complete genome | Apium graveolens | 568 | 663 | 12% | 9e-156 | 91.12% | 263017 | MZ328722.1 |
| Apium graveolens strain L.+ W99B mitochondrion, complete genome | Apium graveolens | 568 | 663 | 12% | 9e-156 | 91.12% | 394073 | MK562756.1 |
| Cicuta virosa chromosome 1 mitochondrion, complete sequence | Cicuta virosa | 547 | 547 | 11% | 1e-149 | 92.53% | 352718 | PQ423759.1 |
| Cuscuta gronovii cytochrome c maturation subunit Fn (ccmFn) gene, complete cds; and trnL pseudogene, co... | Cuscuta gronovii | 544 | 544 | 18% | 1e-148 | 83.41% | 18827 | KP940496.1 |
| Ilex metabaptista mitochondrion, complete genome | Ilex metabaptista | 540 | 995 | 21% | 2e-147 | 90.16% | 529560 | NC_081509.1 |
| Ilex macrocarpa mitochondrion, complete genome | Ilex macrocarpa | 534 | 990 | 21% | 9e-146 | 89.93% | 539461 | NC_082235.1 |

**Figure S7.** The sequence of OG0000602-612 was aligned against the NCBI NT database using BLAST in NCBI.

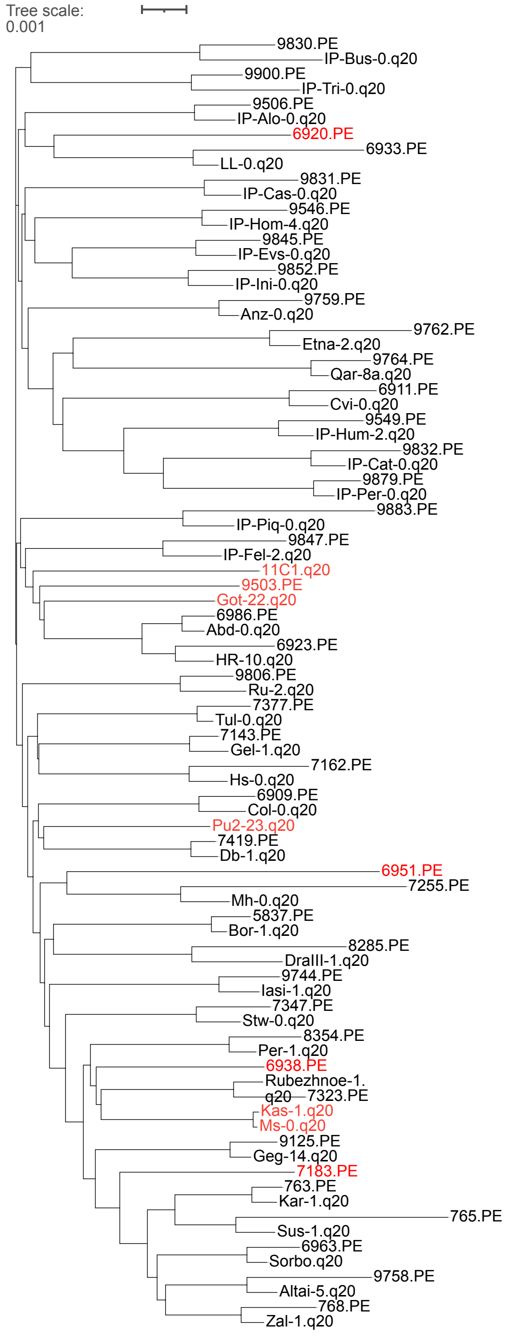


**Figure S8.** Phylogenetic Tree of Short Read and HiFi Read Data. IDs with the "q20" suffix represent HiFi data, while IDs with the "PE" suffix represent short-read data. Accessions with inconsistencies are marked in red and were excluded from the analysis.

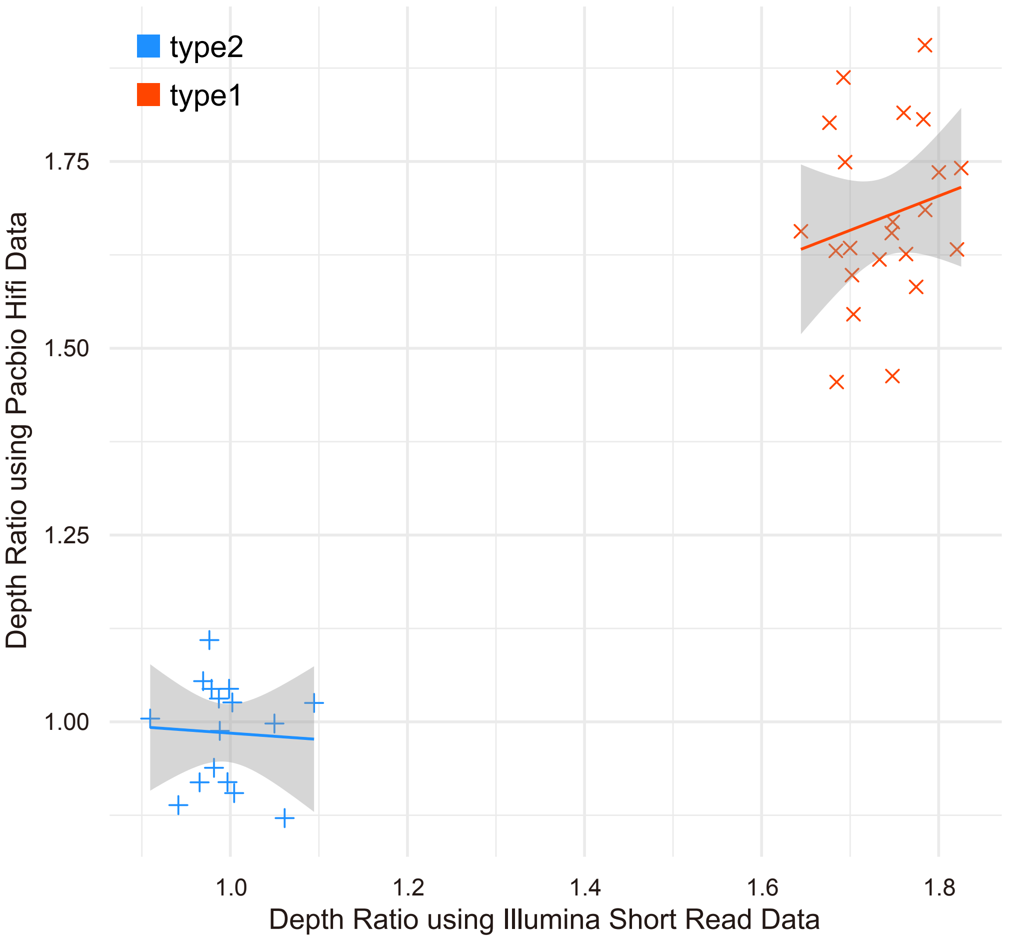


**Figure S9.** Distribution of Depth Ratios Inferred from Short Reads and HiFi Reads. Each dot represents one accession.

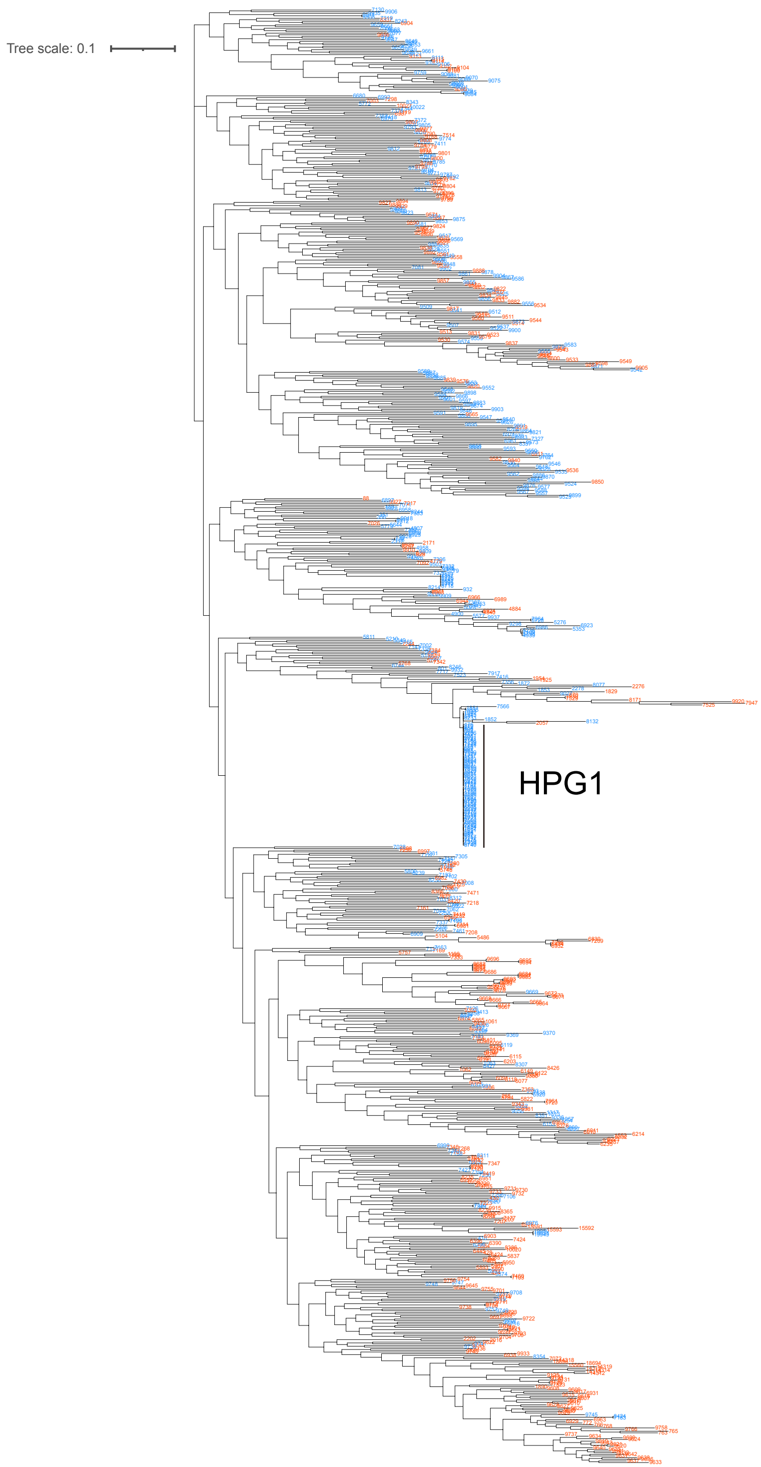


**Figure S10.** Distribution of Type 1 and Type 2 Mitochondria genomes projected onto the phylogenetic tree. Red IDs represent Type 1, while blue IDs represent Type 2.

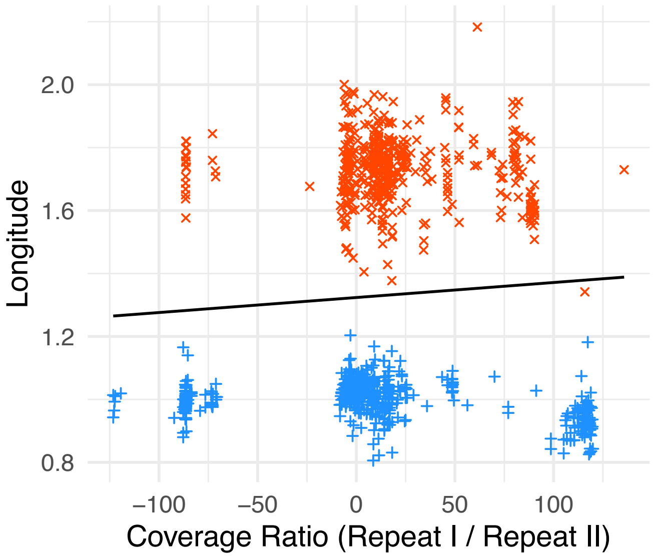


**Figure S11.** Correlation between coverage ratio and longitude.
